# The *Drosophila melanogaster* PIF1 helicase promotes survival during replication stress and processive DNA synthesis during double-strand gap repair

**DOI:** 10.1101/554550

**Authors:** Ece Kocak, Sarah Dykstra, Alexandra Nemeth, Catherine G. Coughlin, Kasey Rodgers, Mitch McVey

## Abstract

PIF1 is a 5’ to 3’ DNA helicase that can unwind double-stranded DNA and disrupt nucleic acid-protein complexes. In *Saccharomyces cerevisiae*, Pif1 plays important roles in mitochondrial and nuclear genome maintenance, telomere length regulation, unwinding of G-quadruplex structures, and DNA synthesis during break-induced replication. Some, but not all, of these functions are shared with other eukaryotes. To gain insight into the evolutionarily conserved functions of PIF1, we created *pif1* null mutants in *Drosophila melanogaster* and assessed their phenotypes throughout development. We found that *pif1* mutant larvae exposed to high concentrations of hydroxyurea, but not other DNA damaging agents, experience reduced survival to adulthood. Embryos lacking PIF1 fail to segregate their chromosomes efficiently during early nuclear divisions, consistent with a defect in DNA replication. Furthermore, loss of the BRCA2 protein, which is required for stabilization of stalled replication forks in metazoans, causes synthetic lethality in third instar larvae lacking either PIF1 or the polymerase delta subunit POL32. Interestingly, *pif1* mutants have a reduced ability to synthesize DNA during repair of a double-stranded gap, but only in the absence of POL32. Together, these results support a model in which Drosophila PIF1 functions with POL32 during times of replication stress but acts independently of POL32 to promote synthesis during double-strand gap repair.

## Introduction

Pif1 family helicases are 5’ to 3’ superfamily 1 helicases that are highly conserved in most eukaryotes and some bacteria and are critical for DNA replication, recombination, and repair (Bochman *et al.* 2010; Chung 2014; Byrd and Raney 2017). Although Pif1 family helicases possess a conserved single-stranded (ss) DNA-dependent helicase domain, their N- and C-terminal domains differ significantly in size and sequence between organisms (Figure S1) (Lahaye *et al.* 1991; Boule and Zakian 2006). Furthermore, the cellular functions of PIF1 homologs in eukaryotes are variably conserved. Because human PIF1 appears to act as a tumor suppressor but is also required for the survival of cancer cells (Gagou *et al.* 2014), a better understanding of its contribution to genome stability in diverse cellular and organismal contexts is needed.

Much of what is currently known about Pif1 structure and function comes from studies in the yeasts *Saccharomyces cerevisiae* and *Schizosaccharomyces pombe*. *S. cerevisiae* possesses two Pif1 orthologs, ScPif1 and ScRrm3, which localize to both the mitochondria and nucleus (Ivessa *et al.* 2000; Bochman *et al.* 2010). ScPif1 was first discovered in a genetic screen where its deficiency resulted in reduced mitochondrial (mt) DNA recombination (Foury and Kolodynski 1983). Further studies revealed that ScPif1 slows down replication progression in mtDNA to prevent double-stranded breaks (DSBs) and cooperates with base excision repair to resolve oxidative mtDNA damage (O’rourke *et al.* 2002; Doudican *et al.* 2005; Cheng *et al.* 2007). Similar to ScPif1, *S. pombe* Pfh1 (PIF1-homolog-1) maintains mtDNA integrity. In addition, both ScPIF1 and SpPfh1 maintain telomeric DNA integrity and normal telomere length (Schulz and Zakian 1994; Mcdonald *et al.* 2014). However, loss of Pfh1 is lethal (Zhou *et al.* 2002; Pinter *et al.* 2008), while deletion of *S. cerevisiae* Pif1 is not, for reasons that are still not understood.

Pif1 appears to play multiple roles during DNA replication. ScPif1 promotes Okazaki fragment maturation during lagging strand synthesis by contributing to the displacement of downstream Okazaki fragments and the production of longer single-strand flaps by increasing DNA polymerase delta (Pol δ) processivity (Rossi *et al.* 2008; Stith *et al.* 2008). SpPfh1 likewise contributes to the formation of single-strand flaps and resolves secondary structures at these flaps to promote their cleavage by the Dna2 nuclease (Tanaka *et al.* 2002; Ryu *et al.* 2004). Pfh1 has additional essential roles in DNA replication, as it interacts with multiple replisome core proteins and is required for fork merging at replication termination sites (Tanaka *et al.* 2002; Steinacher *et al.* 2012). Recently, ScPif1 was also shown to be important for centromere replication in the absence of the Rrm3 protein (Chen *et al.* 2019)

One of PIF1’s most conserved roles is the resolution of G quadruplex (G4) DNA. G4 DNA are stable secondary structures formed by the association of four guanines in G-rich DNA held together by non-canonical Hoogsteen base pairs (Maizels and Gray 2013). The formation of G4 structures *in vivo* can cause problems for DNA replication due to their high thermal stability (Lipps and Rhodes 2009; Bochman *et al.* 2012). Both ScPif1 and SpPfh1 unwind intra- and intermolecular G4 structures *in vitro* (Paeschke *et al.* 2011; Sabouri *et al.* 2014; Wallgren *et al.* 2016). In addition, both helicases can remove stably bound protein complexes from DNA to allow for DNA replication and to prevent G4 formation at lagging strand telomeres (Sabouri *et al.* 2012; Galletto and Tomko 2013; Koc *et al.* 2016). The G4 unwinding activity has been shown to have *in vivo* relevance, as DNA replication progresses more slowly near G4 motifs in Pif1 deficient cells (Paeschke *et al.* 2011; Paeschke *et al.* 2013).

In yeasts, Pif1 plays important roles in DNA repair. *S. cerevisiae pif1* mutants show mild sensitivity to methyl methanesulfonate (MMS), a DNA alkylating agent, and hydroxyurea (HU), a ribonucleotide reductase inhibitor that depletes nucleotide pools and stalls replication forks (Budd *et al.* 2006; Wagner *et al.* 2006). Interestingly, the sensitivity of *S. pombe pfh1* mutants to these chemicals is much greater (Tanaka *et al.* 2002; Pinter *et al.* 2008). ScPif1 co-localizes with the homologous recombination (HR) protein Rad52 to repair foci after gamma irradiation, and SpPfh1 is recruited to DNA damage foci in camptothecin-treated cells.

Recently, ScPif1 was shown to be required during break-induced replication (BIR), a type of HR repair observed at one-ended DNA breaks caused by replication fork collapse or shortening of telomeres (Saini *et al.* 2013; Buzovetsky *et al.* 2017). During BIR, Rad51-mediated invasion of ssDNA into a homologous template creates a migrating bubble, resulting in conservative synthesis of a nascent DNA duplex (Donnianni and Symington 2013; Kramara *et al.* 2018). Efficient BIR depends upon both Pif1 and Pol32, a non-catalytic subunit of Pol δ (Wilson *et al.* 2013). Currently models suggest that ScPif1 promotes Pol δ-dependent DNA synthesis during the bubble migration and may also unwind the newly-synthesized strand to relieve topological stress (Sakofsky and Malkova 2017).

Several recent PIF1 studies have revealed that some of its roles are conserved in metazoans. Human PIF1 localizes to both nuclei and mitochondria. It interacts with the catalytic reverse transcriptase subunit of telomerase and its overexpression results in decreased telomere length (Mateyak and Zakian 2006; Zhang *et al.* 2006; Futami *et al.* 2007; Paeschke *et al.* 2013). Mice lacking PIF1 show a mitochondrial myopathy, suggesting a subtle role in mtDNA maintenance (Bannwarth *et al.* 2016). The expression of mouse and human PIF1 is limited to highly proliferating embryonic stem cells and peaks in late S/G2 phase, consistent with a role in DNA replication (Mateyak and Zakian 2006). In addition, human PIF1 is important for S-phase entry and is thought to allow unperturbed DNA replication by unwinding G4 DNA and stalled replication fork-like substrates (George *et al.* 2009; Sanders 2010).

However, the roles and phenotypes of the yeast Pif1 helicases are not always consistent with those of its metazoan orthologs. For example, mice lacking PIF1 show no visible phenotypes, have unaltered telomere length, and display no chromosomal abnormalities. These observations are consistent with a recent study demonstrating substantial evolutionary divergence between yeast and human PIF1 proteins (Dehghani-Tafti *et al.* 2019) and suggest that Pif1 may play a subtler role in higher eukaryotes or may have redundant functions with other helicases (Snow *et al.* 2007).

To investigate the possible roles of PIF1 in a non-mammalian metazoan, we generated and characterized a *pif1* null mutant in *Drosophila melanogaster*. Here, we show that *pif1* mutants are sensitive to hydroxyurea and exhibit reduced embryo viability associated with nuclear fallout, chromosome clumping, anaphase bridges, and asynchronous mitotic divisions. Both phenotypes suggest that PIF1 plays an important function during periods of replication stress, including that encountered during embryogenesis. In addition, we demonstrate a role for Drosophila PIF1 in promoting long-range DNA synthesis during HR repair, specifically in the absence of POL32. Finally, we identify a specific genetic interaction between PIF1 and BRCA2, which functions both in double-strand break repair and replication fork protection in mammals. Together, our findings suggest that Drosophila PIF1 shares DNA replication and double-strand break related functions of its yeast counterparts and may play an additional role in the protection of stalled or regressed replication forks.

## Materials and Methods

### Drosophila stocks and mutants

Drosophila stocks were maintained on standard cornmeal agar medium at 25°C. The *pif1^167^* null mutant was created via imprecise excision of a *P*-element, (*P{EPgy2}Pif1^EY10295^*, Bloomington Stock 17658) located in the 5’ untranslated region of *CG3238* (Alexander *et al.* 2016). The deletion removes nucleotides 64-1759 (relative to the transcription start site) and inserts the sequence CTGTTATTTCATCATG at the deletion breakpoint.

Other mutants used in this study include *pol32^L2^*, which removes all but the first 42 nucleotides of the POL32 coding sequence (Kane *et al.* 2012); *brca2^47^*, which deletes the first 2169 bp of the 3417 bp BRCA2 coding sequence (Thomas *et al.* 2013); *brca2^KO^*, which deletes the first 3321 bp of the BRCA2 coding sequence (Klovstad *et al.* 2008); *rad51^057^* (A205V, a null mutation), (Staeva-Vieira *et al.* 2003); and *mus81^Nhe^*, a 16nt insertion in the coding region (Trowbridge *et al.* 2007). All double and triple mutants used in this study were created by standard genetic crosses and verified by PCR or Sanger sequencing. The H2Av-EGFP stock was obtained from Bloomington Drosophila Stock Center (Clarkson and Saint 1999).

### Mutagen sensitivity assays

Sets of five virgin females and three males heterozygous for the *pif1^167^* mutation were mated in vials containing standard cornmeal agar medium. Females were allowed to lay eggs for 3 days before being transferred into a second vial to lay for an additional 3 days. The first set of vials served as the controls and was treated with 250 μL of water 1 day after the transfer of parents. The second set of vials was treated with 250 μL of the mutagen. Upon eclosion, adults were counted in each vial and the percentage of viable homozygotes in each vial was determined relative to the total number of adult progeny. Relative survival was calculated as (% viable homozygotes in the mutagen-treated vials)/(% viable homozygotes in control vials) for each trial. Statistical significance was determined using paired t-tests between untreated and treated vials.

### Hatching frequency assay

Hatching frequency was measured by mating 30-40 virgin female flies to 15-20 male flies for each genotype. The mating population was placed in a single vial with dry yeast overnight at 25°C to acclimate them for mating and laying. After 24 hours, the flies were transferred to an egg collection chamber consisting of a 100 mL Tri-Stir beaker with small holes in the bottom capped with a yeasted grape juice agar plate. The flies were left to mate and lay eggs for approximately 4-5 hours or until each plate had about 100-150 eggs present on the grape agar. Hatching frequency was visually scored using a stereomicroscope 48 hours after egg collection for wild type or 72 hours after collection for *pif1^167^* mutants. Three trials of 3 plates for each genotype’s progeny were scored. Statistical significance was determined using unpaired t-tests between the three mating groups.

### Embryo collection and DAPI staining

Thirty *pif1^167^* homozygous mutants (3:1 female to male ratio) were collected and placed in a vial with dry yeast overnight at 25°C. The next day, the population was placed in an egg collection chamber and returned to 25°C to lay. Grape agar plates containing newly laid embryos were collected and changed every 2 hours. The embryos were collected from each grape agar plate and dechorionated in 50% bleach for 2 minutes. Embryos were then fixed with a mixture of 1:1 heptane:37% formaldehyde for 20 minutes, stained with 1 μg/mL DAPI in 0.1% Triton-X PBS for 3 minutes, and mounted onto a glass slide with Vectashield mounting media. The embryos were covered with a 18×18mm No 1 coverslip which was sealed with clear nail polish. Embryos were visualized on a Zeiss Axio Z-stepping microscope and fluorescent images were acquired with a Zeiss Axiocam 503 mono camera. Quantification of the frequency of nuclear defects for DAPI-stained *pif1* and wild-type embryos was performed on deconvolved images. Statistical significance was determined using unpaired t-tests.

### Time-lapse microscopy

About 100 wild-type H2Av–EGFP and *pif1^167^*–H2Av–EGFP homozygous flies (3:1 female to male ratio) were collected and placed in separate bottles with dry yeast overnight at 25°C. After 24 hours, each population was placed in an egg collection chamber and returned to 25°C to lay. Grape agar plates containing newly laid embryos were removed every hour. Embryos were then collected and manually dechorionated. Dechorionated embryos were transferred onto a slide containing Halocarbon Oil 700. The embryos were covered with a 18×18mm No 1 coverslip, while maintaining enough distance between the slide and the coverslip by use of 4 layers of double-sided tape. Embryos were visualized on a Zeiss Axio Z-stepping microscope and fluorescent images were acquired with a Zeiss Axiocam 503 mono camera every 20 seconds.

### Nondisjunction assay

Meiotic nondisjunction of the *X* chromosomes was measured by crossing *y w* or *y w; pif1^167^* females to *y w^a^ Ste^1^* /*Dp(1;Y)B^S^ y^+^* males. The duplication on the *Y* chromosome carries a dominant mutation causing bar-shaped eyes. Fertilization of eggs produced through normal meiotic disjunction produces females with non-Bar eyes and yellow bodies and males with Bar eyes and wild-type bodies. Fertilization of diplo-*X* ova created through nondisjunction results in *XXY* female progeny with Bar eyes and normal-colored bodies (and non-viable *XXX* progeny). Fertilization of nullo-*X* ova results in *XO* males with non-Bar eyes and yellow bodies (and non-viable *YO* progeny). *X* nondisjunction was calculated as the percentage of progeny that arose from nondisjunction (Bar females and non-Bar males), after correcting for loss of half of the diplo-*X* ova and half of the nullo-*X* ova.

### *P{w^a^}* site-specific gap repair assay

*P{w^a^}* is a 14-kilobase *P* element containing the *white* gene, interrupted by a *copia* retrotransposon. In this assay, it is inserted in the essential gene *scalloped* on the *X* chromosome. A third chromosome transposase source (*P{ry^+^, Δ2-3}99B*, Bloomington stock 406) was used to excise *P{w^a^}*. Following excision of *P{w^a^}*, double-strand gap repair events occurring in the male pre-meiotic germline were recovered and analyzed as described previously (Mcvey 2010).

Single males possessing both *P{w^a^}* and transposase were mated to females homozygous for *P{w^a^}* in individual vials and repair products were recovered in female progeny. Repair events with full HR repair, involving synthesis through the *copia* long terminal repeats (LTRs) are recovered in red-eyed females, while events with aborted HR repair followed by end joining are recovered in yellow-eyed females. Individual progeny were scored for eye color and statistical significance was determined using the Mann-Whitney test, with each vial considered an independent experiment. Genomic DNA from flies possessing independent repair events (one per vial) was recovered (Gloor *et al.* 1993) and PCR with primers tiled across *P{w^a^}* were used to estimate the extent of repair synthesis. Statistical significance for repair synthesis tract lengths was determined with Fisher’s exact test.

### Life stage-specific synthetic lethality

10-15 virgin females and 5 males both heterozygous for the mutation of interest *in trans* to a GFP-bearing balancer chromosome were placed in a vial with dry yeast overnight at 25°C. After 24 hours, the flies were transferred to an egg collection chamber and females were allowed to lay eggs for 5 hours. The resulting progeny were observed daily from the onset of first instar larvae to eclosion. Heterozygotes and homozygotes were scored by presence or absence of GFP, respectively. Time of synthetic lethality was determined by the life stage at which no homozygotes were observed.

### Data and reagent policy

Fly stocks are available upon request. Supplemental files are available at FigShare. File S1 contains supplemental figures. File S2 contains time-lapse videos of embryonic nuclear divisions in wild-type and *pif1* mutant embryos.

## Results

### *pif1* mutants are sensitive to agents that induce replication stress

We identified a *Drosophila melanogaster* PIF1 ortholog encoded by *CG3238*, based on sequence similarity with PIF1 proteins from other eukaryotes (Figure S1). Like other PIF1 family members, Drosophila PIF1 possesses a P-loop containing triphosphate hydrolase, indicative of ATP-dependent helicase activity, and the 21 amino acid Pif1 signature motif (Bochman *et al.* 2010). However, unlike the yeast Pif1 proteins, Drosophila PIF1 does not appear to have a mitochondrial targeting sequence, suggesting that it may not function in mitochondrial maintenance. To investigate the roles of PIF1 in Drosophila, we used the previously generated *pif1^167^* mutant described in (Alexander *et al.* 2016). This mutant was created via imprecise excision of a *P* element (*EY10295*) located in the 5’ untranslated region of *CG3238*, which resulted in the removal of the majority of the helicase domain. Flies homozygous for this deletion were viable and apparently healthy, although females displayed greatly reduced fertility due to maternal effect lethality (see next section).

*S. pombe pfh1* mutants are extremely sensitive to compounds that induce replication stress, including the alkylating agent methyl methanesulfonate (MMS) and the ribonucleotide reductase inhibitor hydroxyurea (HU) (Tanaka *et al.* 2002), while *S. cerevisiae pif1* mutants exhibit milder sensitivity to these agents (Budd *et al.* 2006). To investigate the roles of Drosophila PIF1 in responding to various types of DNA damage, we treated *pif1^167^* heterozygous and homozygous mutant larvae with increasing concentrations of different mutagens and measured the survival of these larvae into adulthood relative to untreated siblings. The mutagen concentrations were selected based on our previous findings that other DNA repair mutants are sensitive at these doses (Kane *et al.* 2012; Thomas *et al.* 2013; Beagan *et al.* 2017). Notably, as the dose of each mutagen increased, the total number of flies (both heterozygotes and homozygotes) that survived decreased, indicating the efficacy of the treatments (Figure S2). The homozygous mutants were not sensitive to the highest concentrations of nitrogen mustard (which induces intra- and interstrand DNA crosslinks), topotecan (a topoisomerase I inhibitor that creates one-ended DSBs following DNA replication), or paraquat (induces oxidative damage), and showed only a mild sensitivity to the alkylating agent methyl methanesulfonate and the radiomimetic drug bleomycin (Figure 1A). Interestingly, *pif1* homozygous mutants showed moderate sensitivity when treated with a high dose of hydroxyurea, which can stall replication forks and eventually cause fork collapse (Figure 1A). Further testing showed that loss of PIF1 causes a dose-dependent increase in hydroxyurea sensitivity (Figure 1B). Together, these results suggest that Drosophila PIF1 does not play a major role in responding to oxidative stress or bulky DNA lesions, unlike its yeast orthologs, but is important for survival during conditions of replication stress.

**Figure 1.**
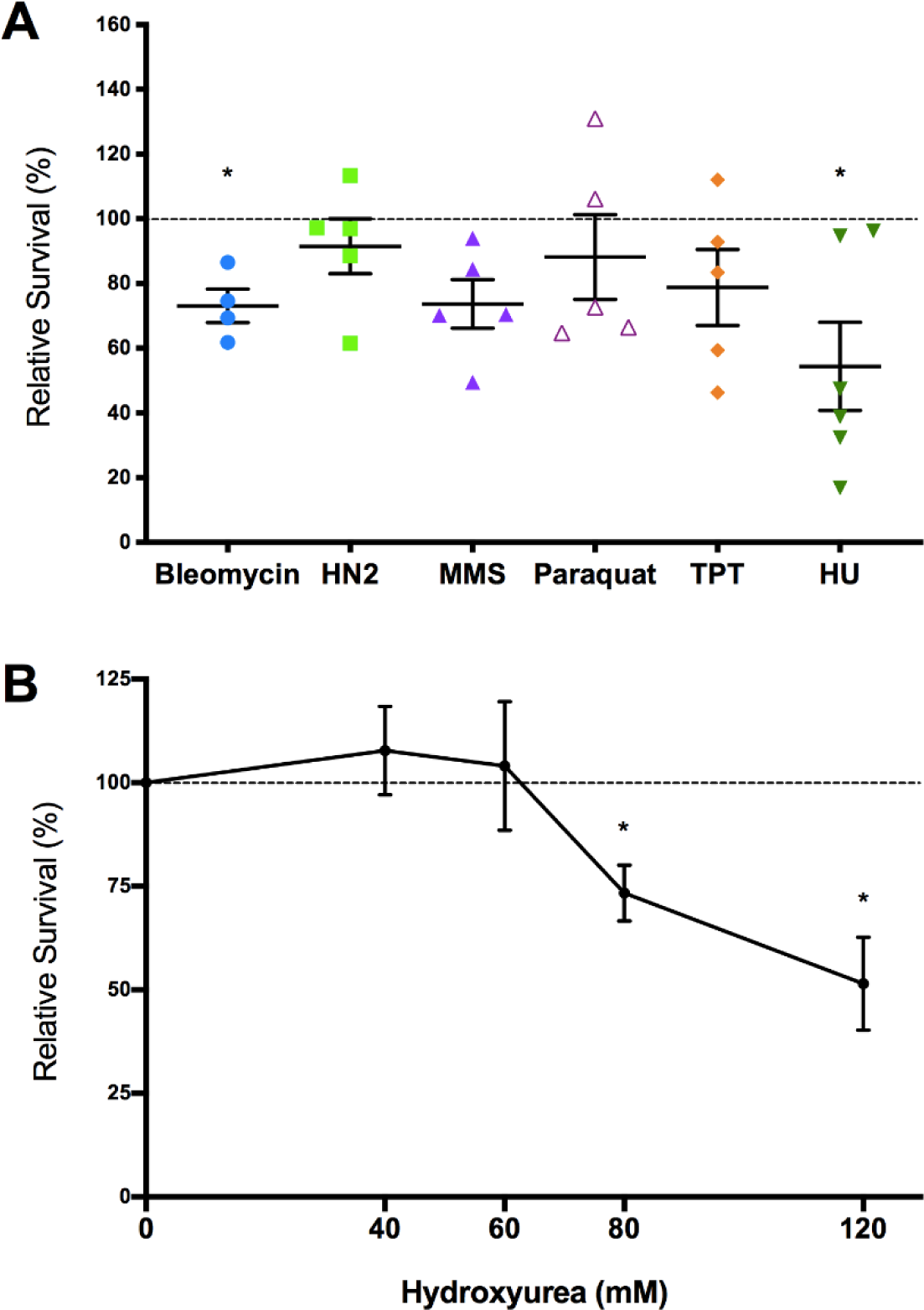
Drosophila PIF1 mutants are mildly sensitive to hydroxyurea. (A) Flies heterozygous for the *pif^167^* null mutation were mated in vials for three days and then transferred to new vials for two days. The first set of control vials was treated with water and the second set was treated with solutions of 25 uM bleomycin, 0.01% nitrogen mustard (HN2), 0.12% methane methylsulfonate (MMS), 1 mM paraquat, 50 uM topotecan (TPT), or 120 mM hydroxyurea (HU). Relative survival for each pair of vials was calculated as the ratio of the percentages of homozygous mutant flies in treated vs. untreated vials. Each point represents a biological replicate (vial pair), with the mean and SEM shown. The dotted line shows 100% relative survival. * indicates P < 0.05 in paired t-tests between control and mutagen treated vials. (B) Flies heterozygous for the *pif1^167^* mutation were mated in vials and treated with water of increasing concentrations of hydroxyurea. The mean and SEM of at least six biological replicates is shown for each HU concentration. Dotted line indicates 100% relative survival. * indicates P < 0.05 in paired t-tests between unexposed and exposed vials.

### PIF1 is required for rapid nuclear divisions in early embryogenesis

During our initial characterization of the *pif1^167^* allele, we found that we were unable to maintain a stock of homozygous *pif1* mutants, as homozygous mutant females failed to produce viable progeny. Because *pif1* mutants are preferentially sensitive to agents that induce replication stress, we hypothesized that this defect might be due to embryonic lethality resulting from replication defects during early development. Indeed, embryos obtained from *pif1* mutant mothers had a significantly decreased hatching frequency (13.6%) relative to wild-type embryos (87.1%, Figure 2). The small percentage of *pif1* mutant embryos that did hatch developed more slowly than wild-type embryos, frequently taking up to 48 hours to emerge as first-instar larvae as opposed to 24 hours for wild type.

**Figure 2.**
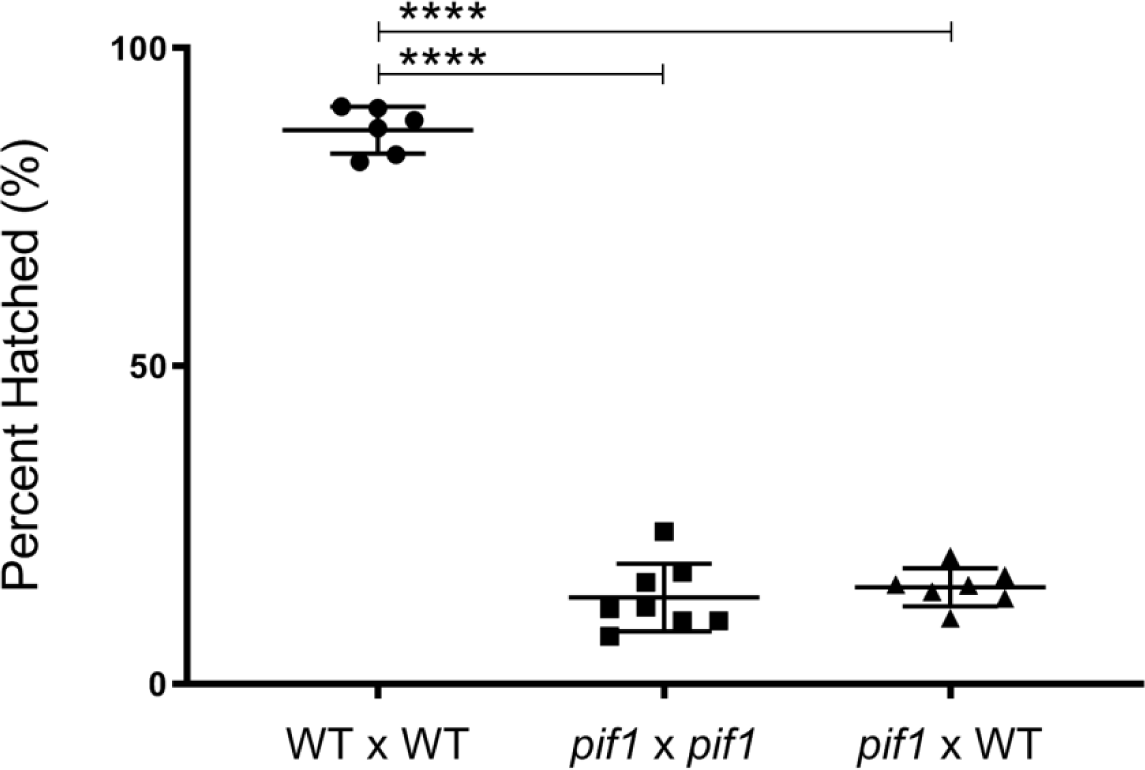
Severely decreased hatching frequency in *pif1* mutant flies is due to an early embryonic function of PIF1. Embryos were collected from wild type x wild type, *pif1^167^ x pif1^167^*, and *pif1^167^*(female) x *w^1118^* (male) matings, maintained at 25°C for 72 hours after collection, and scored for hatching. For each mating, three independent trials with at least two egg collections per trial were performed, with at least 550 embryos scored for each trial. Each point corresponds to the percentage of eggs hatched per plate. **** corresponds to P <0.0001 in unpaired t-tests between the three mating groups.

In Drosophila embryogenesis, paternal gene expression of most genes is initiated about two hours after fertilization. To test whether the contribution of the functional paternal PIF1 could rescue the severe defect in hatching frequency seen in the *pif1* mutant embryos, we mated *pif1* homozygous females with wild-type males. The average hatching frequency of the resultant *pif1* heterozygous embryos was 15.2% and was not significantly different from the *pif1* mutant females mated to *pif1* mutant males. This inability of paternally-derived PIF1 to rescue the *pif1* egg-hatching defects suggests that the crucial function of PIF1 occurs during the first two hours of egg development.

During the early stages of Drosophila embryo development, alternating S and M phases occur every five to six minutes to produce approximately 6000 syncytial nuclei in a common cytoplasm (Zalokar and Erk 1976). Nuclei that fail to complete replication and/or experience aberrant mitotic chromosome segregation are removed through an active process termed nuclear fallout (Sullivan and Theurkauf 1995). To determine if DNA replication and subsequent nuclear division is affected in the absence of PIF1, we fixed wild-type and *pif1* homozygous mutant embryos and stained them with the fluorescent DNA marker DAPI to visualize the syncytial nuclei within 1-2 hours after fertilization. Wild-type embryos undergoing normal syncytial divisions exhibited an even spatial pattern of nuclei (Figure 3A). In contrast, we observed multiple nuclear defects in *pif1* mutants undergoing embryogenesis, including gaps in the typically uniform monolayer of nuclei, abnormal chromosome clumping, chromosome fragmentation, anaphase bridges, and asynchronous mitotic divisions (Figure 3B). Quantification of these defects showed that nearly all *pif1* embryos exhibit nuclear fallout and chromosome clumping, while almost half show evidence of anaphase bridges (Figure 3C). These phenotypes were also present in very early nuclear division cycles (Figure S3), suggesting that the requirement for PIF1 begins at the earliest stages of embryogenesis.

**Figure 3.**
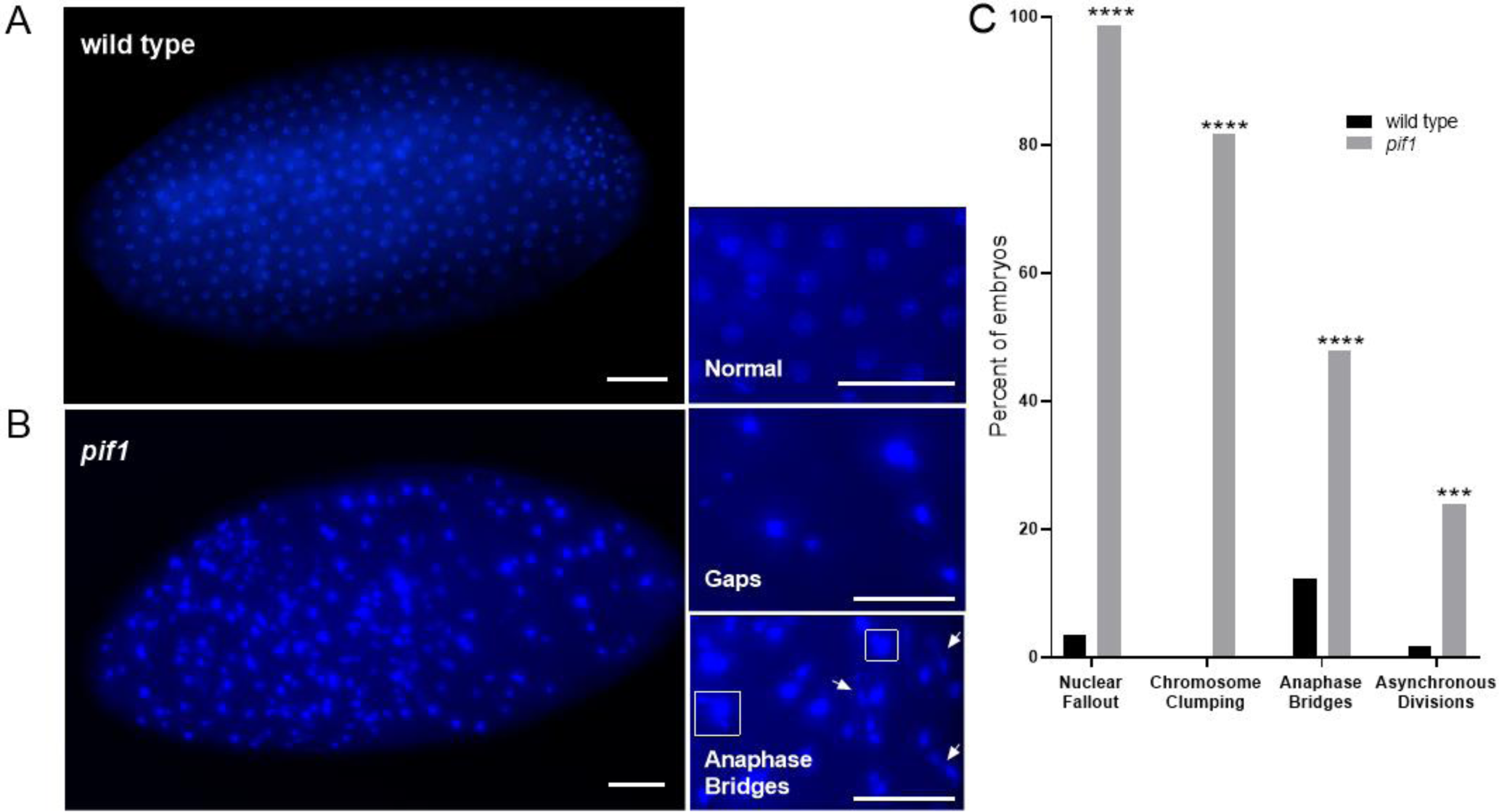
*pif1* mutant embryos exhibit severe chromosome segregation defects. Embryos from both wild-type and *pif1^167^* mutant flies were collected after 1-2 hours of development at 25C, stained with DAPI, and visualized at 40X magnification using florescence microscopy. (A) Representative wild-type embryo with normal nuclear patterning. Bar = 50 μm (B) Representative *pif1* mutant embryo showing gaps within the normally uniform nuclear monolayer, anaphase bridges, and nuclear clumping. Bar = 50 μm. (C) Quantification of the frequency of nuclear defects seen in DAPI-stained wild-type and *pif1^167^* embryos. wild type n = 57; *pif1^167^*n = 71. **** P < 0.0001, *** P < 0.001 in unpaired t-tests.

To further characterize the developmental defects in *pif1* mutants, we employed real-time visualization of developing wild-type and *pif1* embryos. We introduced constructs encoding the histone H2A variant fused to enhanced green fluorescent protein (H2Av–EGFP) into wild-type and *pif1* mutant backgrounds and followed the nuclei through multiple mitoses. We observed nuclear fallout, anaphase bridging, and chromosome clumping in *pif1*, but not wild-type embryos, consistent with the results from the DAPI-stained fixed embryos (Supplemental Videos 1 and 2). Because failure to complete DNA replication during early embryogenesis can lead to chromosome segregation failures similar to what we observed in the *pif1* mutants (Sibon *et al.* 2000), we conclude that PIF1 is likely acting during DNA replication during early Drosophila development.

In Drosophila, defects in meiotic recombination often cause an increase in chromosome non-disjunction (Hughes *et al.* 2018). Thus, the maternal effect lethality observed in *pif1* mutants could also be caused by defective meiotic recombination. To test this, we crossed wild type and *pif1* mutants to males possessing a *Y* chromosome with a dominant marker *B^S^* (see materials and methods). While the frequency of nondisjunction for the *pif1* mutants was higher than the wild-type frequency (Figure S4), the increase was not statistically significant due to the small number of adults that we were able to recover from *pif1* mutant mothers. Thus, we conclude the maternal effect lethality observed in *pif1* mutants is largely caused by an inability to carry out efficient DNA replication, although we cannot rule out the possibility that meiotic recombination defects may also play a role.

### PIF1 promotes long-range DNA synthesis during homologous recombination repair

While *S. cerevisiae* Pif1 plays a minimal role during gene conversion via homologous recombination, it is essential for break-induced replication (BIR), a type of homologous recombination repair that requires large amounts of synthesis and proceeds through migration of a mobile D-loop (Saini *et al.* 2013; Wilson *et al.* 2013). The POL32 subunit of Pol δ is also required for BIR (Wilson *et al.* 2013) and has been shown to promote extensive synthesis during gap repair in Drosophila (Kane *et al.* 2012). To determine whether there is a similar requirement for PIF1 during gap repair via homologous recombination, we used the well-characterized *P{w^a^}* assay (Mcvey 2010). This assay uses a *X*-linked *P* element containing an interrupted *white* gene (Figure 4A). The excision of *P{w^a^}* from one sister chromatid in the presence of *P*-transposase generates a 14-kb gap that can be repaired via HR using an intact sister chromatid as a template for repair synthesis. Synthesis often terminates prematurely and repair can be completed by end joining.

**Figure 4.**
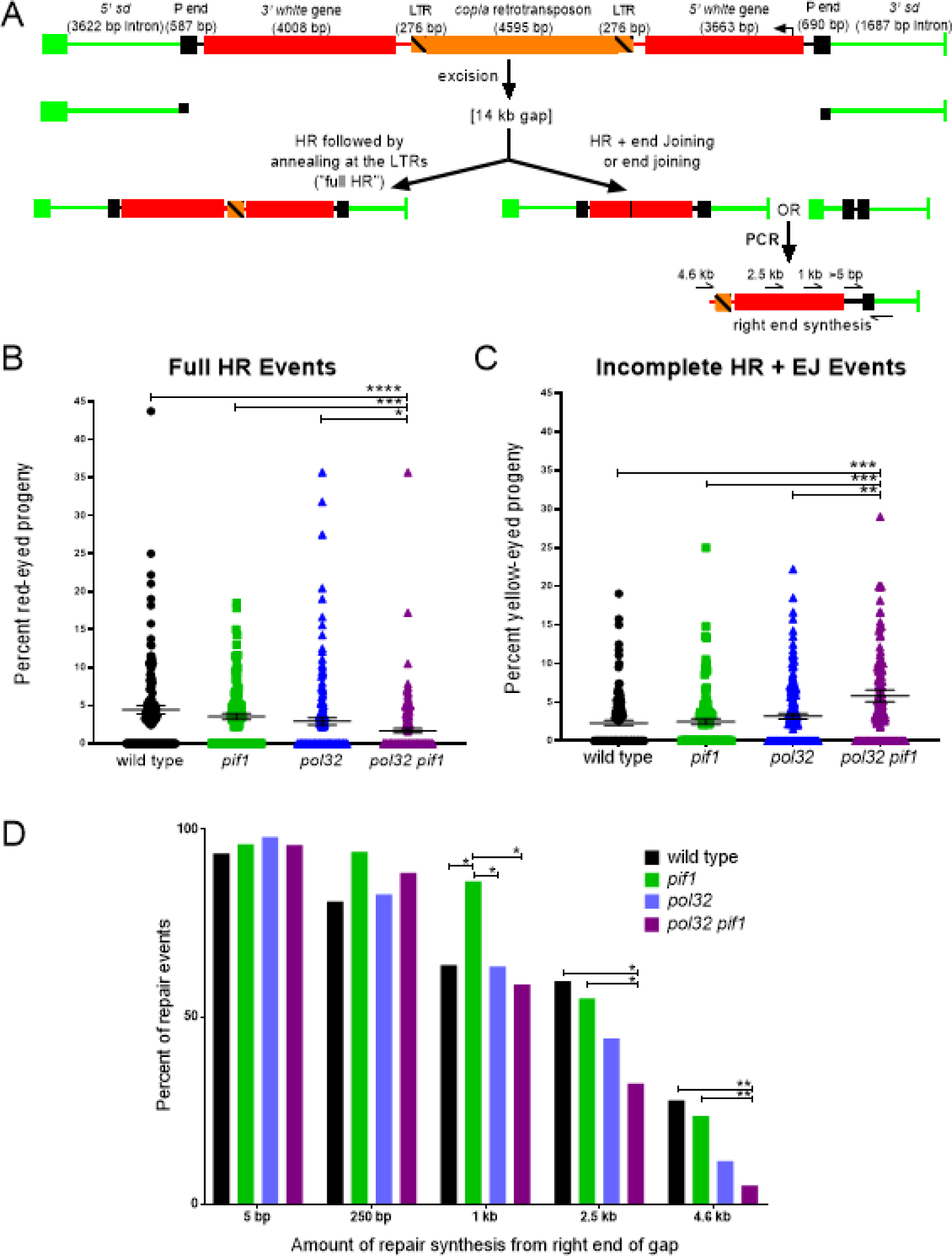
PIF1 and POL32 independently promote DNA synthesis during HR gap repair. (A) The *P{w^a^}* site-specific gap repair assay. A *P* element bearing a *copia* retrotransposon in an intron of the *white* gene is inserted in the essential gene *scalloped* (*sd*). Expression of *P* transposase in *P{w^a^}*-bearing males results in a 14-kb gap with 17 nucleotide non-complementary overhangs relative to the uncut sister chromatid. Repair events from the male pre-meiotic germline are recovered in females in the next generation *in trans* to an intact *P{w^a^}*. Recovery of repair events involving full synthesis of *white*, followed by annealing at the *copia* LTRs, creates female progeny with red eyes. Recovery of repair events involving partial synthesis of *white* followed by end joining, or just end joining alone, creates female progeny with yellow eyes. The amount of repair synthesis can be estimated by PCR. (B&C) *pol32 pif1* mutants have decreased full HR repair and increased incomplete HR + EJ repair relative to either single mutant. Number of independent vials scored: wild type = 126; *pol32* = 153; *pif1* = 111; *pol32 pif1* = 156. Error bars represent standard error of the mean. **** P < 0.0001, *** P < 0.001, ** P < 0.002, * P < 0.05 in Mann-Whitney tests between the four genotypes. (D) Repair synthesis during HR is significantly decreased at distances ≥ 2.5 kb in *pol32 pif1* mutants relative to wild type and either single mutant. Each bar represents the percentage of events with at least the indicated amount of synthesis, measured by PCR. Number of independent repair events tested: wild type = 47; *pif1* = 51; *pol32* = 52; *pol32 pif1* = 121. * P < 0.05, ** P < 0.01 in Fisher’s exact tests between the four genotypes.

We found that *pif1* mutants had a slight reduction in the number of full homologous recombination (full HR) events compared to wild type. However, they did not show any changes in the frequency of incomplete HR events that terminated via end joining (HR + EJ; Figures 4B and C). *pol32* mutants, which showed more of a decrease in full HR events than *pif1* mutants, also did not have a significant change in HR + EJ events compared to wild type, consistent with previous findings (Kane *et al.* 2012). Since PIF1 acts with POL32 to promote BIR and long-tract gene conversion in yeast (Lydeard *et al.* 2007; Wilson *et al.* 2013), we tested the ability of *pol32 pif1* double mutants to synthesize long stretches of DNA in the *P{w^a^}* assay. Strikingly, the percentage of full HR events in *pol32 pif1* mutants was significantly decreased compared to either single mutant or wild type, and the percentage of incomplete HR + EJ events was significantly increased (Figures 4B and C).

To quantify the amount of repair synthesis that took place in the incomplete HR products prior to end joining, we utilized a series of PCRs, focusing on right-side synthesis. Loss of POL32 resulted in decreased repair synthesis, with significant effects at large distances 4.6 kb from the right end of the double-strand gap (Figure 4D). In contrast, *pif1* mutants had slightly increased repair synthesis at distances of 250 bp-1 kb and no defects at large distances. Interestingly, the amount of repair synthesis was significantly decreased in *pol32 pif1* mutants relative to either single mutant at distances greater than 2.5 kb. Together, these results suggest that PIF1 does not play a major role in promoting HR requiring extensive synthesis unless POL32 is also missing, in contrast to what is observed during BIR in budding yeast. However, in the absence of POL32, PIF1 becomes important for long-distance synthesis during HR repair.

We previously observed that in the absence of both POL32 and REV3, the catalytic subunit of translesion synthesis polymerase zeta (Pol ζ), there is a significant decrease in full HR and an increase in incomplete HR + EJ events compared to wild type or either single mutant (Kane *et al.* 2012). These results suggested that both Pol δ and Pol ζ act independently to synthesize DNA during *P{w^a^}* gap repair. Given our finding that Drosophila PIF1 becomes important for long-distance synthesis in the absence of POL32, we investigated whether loss of PIF1 would also result in a further reduction in repair synthesis in a Pol ζ mutant background. Interestingly, we found that *pif1 rev3* double mutants do not show a difference in the frequency of full HR or incomplete HR + EJ events relative to *pif1* single mutants, and repair synthesis is not decreased compared to wild-type gap repair events (Figure 5). This differential requirement for POL32 and PIF1 in the absence of Pol ζ provides further evidence that these two proteins are likely promoting long-distance synthesis during HR via different mechanisms.

**Figure 5.**
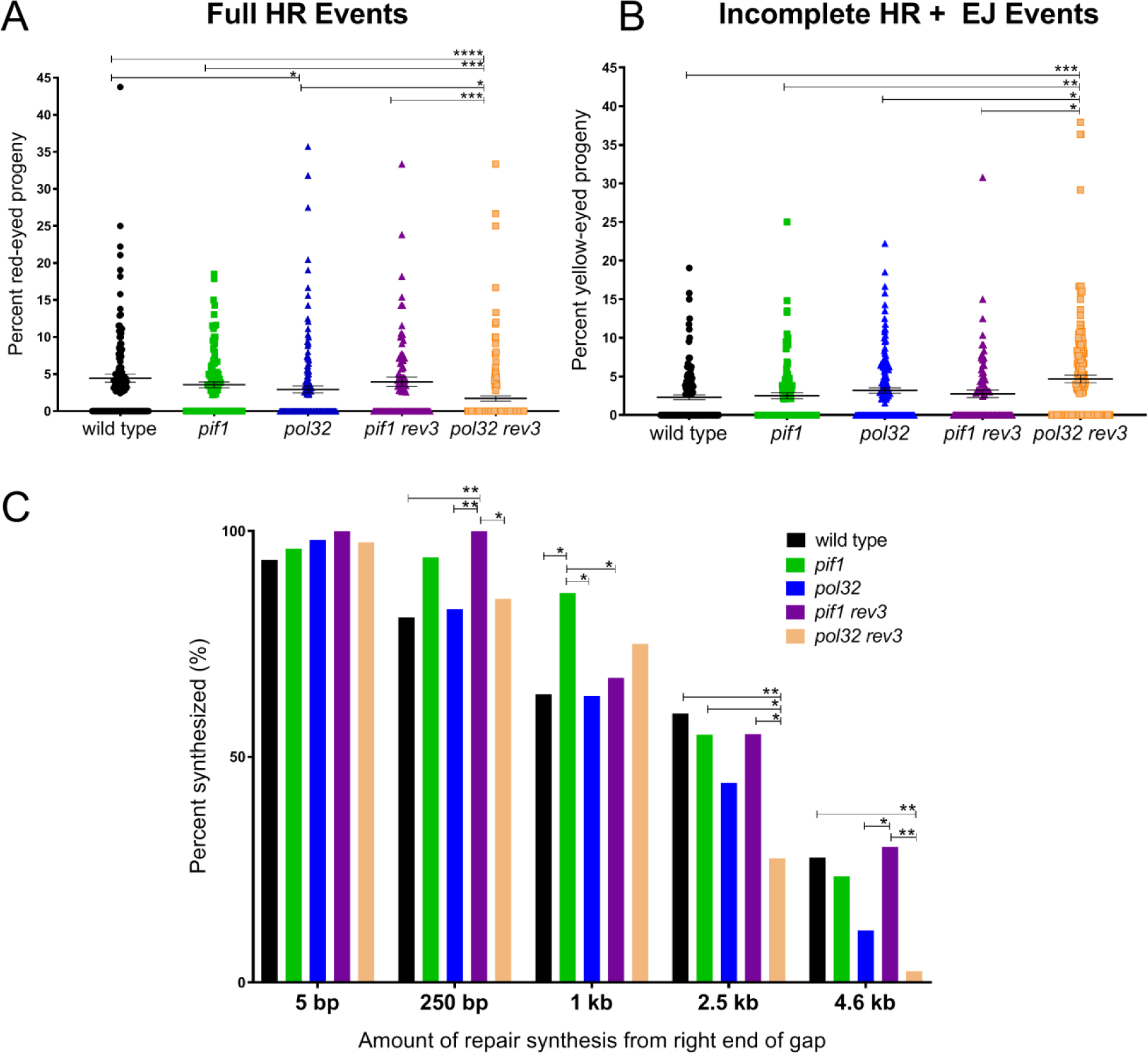
Loss of POL32, but not PIF1, further decreases DNA synthesis during gap repair in *pol zeta* mutants. (A&B) *pol32 rev3*, but not *pif1 rev3* double mutants have decreased full HR and increased incomplete HR + EJ events compared to wild type, *pol32*, or *pif1* single mutants. The wild type, *pif1* and *pol32* data are replicated from Fig. 4. Number of independent vials scored: wild type = 126; *pol32* = 153; *pif1* = 111; *pol32 rev3* = 173; *pif1 rev3* = 86. Error bars represent standard error of the mean. **** P < 0.0001, *** P < 0.001, ** P < 0.002, * P < 0.05 in Mann-Whitney tests between the five genotypes. (C) Repair synthesis is decreased in *pol32 rev3*, but not *pif1 rev3* double mutants relative to wild type, *pif1*, and *pol32* single mutants. Each bar represents the percentage of events with at least the indicated amount of synthesis. Number of independent repair events tested: wild type = 47; *pif1* = 51; *pol32* = 52; *pif1 rev3* = 40; *pol32 rev3* = 40. * P < 0.05, ** P < 0.01 in Fisher’s exact tests between the five genotypes.

### PIF1 and POL32 are essential for adult viability in the absence of BRCA2, but not RAD51

The *P{w^a^}* assay can also be utilized in a *brca2* mutant background to assess how double-strand break repair proceeds in the absence of HR (Mcvey *et al.* 2004; Klovstad *et al.* 2008; Thomas *et al.* 2013). To this end, we generated five independent *pif1 brca2* heterozygous double mutant stocks through meiotic recombination, utilizing two different *brca2* null alleles. Strikingly, we found that *pif1 brca2* homozygous double mutants were unable to survive past late larval stages, precluding their analysis using the *P{w^a^}* assay (Table 1).

**Table 1:** Synthetic lethal relationships for *pif1* and *pol32* mutants

| **Genotype** | **Lethality?** |
| --- | --- |
| *pif1*<sup>167</sup> *brca2*<sup>47</sup> | lethal (3<sup>rd</sup> instar) |
| *pif1*<sup>167</sup> *brca2*<sup>KO</sup> | lethal (3<sup>rd</sup> instar) |
| *pol32*<sup>L2</sup> *brca2*<sup>47</sup> | lethal (3<sup>rd</sup> instar) |
| *pol32*<sup>L2</sup> *brca2*<sup>KO</sup> | lethal (3<sup>rd</sup> instar) |
| *pif1*<sup>167</sup> *pol32*<sup>L2</sup> *brca2*<sup>47</sup> | lethal (3<sup>rd</sup> instar) |
| *pif1*<sup>167</sup> *pol32*<sup>L2</sup> *brca2*<sup>KO</sup> | lethal (3<sup>rd</sup> instar) |
| *pif1*<sup>167</sup> *rad51*<sup>057</sup> | viable adults |
| *pol32*<sup>L2</sup> *rad51*<sup>057</sup> | viable adults |
| *pif1*<sup>167</sup> *brca2*<sup>KO</sup> *rad51*<sup>057</sup> | lethal (3<sup>rd</sup> instar) |
| *pol32*<sup>L2</sup> *brca2*<sup>KO</sup> *rad51*<sup>057</sup> | lethal (3<sup>rd</sup> instar) |
| *pif1*<sup>167</sup> *pol32*<sup>L2</sup> *brca2*<sup>KO</sup> *rad51*<sup>057</sup> | lethal (3<sup>rd</sup> instar) |
| *pif1*<sup>167</sup> *mus81*<sup>Nhe</sup> | viable adults |
| *pol32*<sup>L2</sup> *mus81*<sup>Nhe</sup> | viable adults |
| *pif1*<sup>167</sup> *brca2*<sup>KO</sup> *mus81*<sup>Nhe</sup> | lethal (3<sup>rd</sup> instar) |
| *pol32*<sup>L2</sup> *brca2*<sup>KO</sup> *mus81*<sup>Nhe</sup> | lethal (3<sup>rd</sup> instar) |
| *pif1*<sup>167</sup> *pol32*<sup>L2</sup> *brca2*<sup>KO</sup> *mus81*<sup>Nhe</sup> | lethal (3<sup>rd</sup> instar) |

Because Pol32 acts alongside Pif1 in budding yeast to promote BIR and Okazaki fragment maturation, we tested whether removing POL32 in the absence of HR would also prevent survival to adulthood. Indeed, loss of both POL32 and BRCA2 in two independently derived double mutants resulted in third-instar larval synthetic lethality (Table 1). Similarly, *pol32 pif1 brca2* triple mutants died during late larval stages, suggesting that PIF1 and POL32 act together to promote a process which is required for development past late larval stages in the absence of BRCA2.

In mammals, BRCA2 is required to load RAD51 onto single-stranded DNA during homologous recombination repair of double-strand breaks (Yang *et al.* 2005). It also serves to stabilize stalled replication forks, independent of its function in recombination (Schlacher *et al.* 2011). To determine which of these roles might relate to the lethality observed in *pif1 brca2* mutants, we attempted to create *pif1 rad51* mutants, which like *brca2* mutants should be unable to initiate recombination. Strikingly, we were able to recover the expected Mendelian numbers of viable *pif1 rad51* mutant adults (Table 1 and Figure S5). Similarly, we obtained the expected numbers of *pol32 rad51* adult flies (Figure S5). Based on these data, we conclude that PIF1 and POL32 are not required for adult viability in the absence of homologous recombination.

In mammals, the MUS81 protein acts in a replication fork rescue pathway, cleaving regressed fork that have a single-stranded DNA tail (Hanada *et al.* 2007; Lemacon *et al.* 2017). Following MUS81 cleavage, synthesis is resumed in a POL32-dependent manner. To test whether a similar pathway might operate in Drosophila, we attempted to create flies lacking both MUS81 and PIF1 or POL32. We observed no synthetic lethal interactions between these mutations (Table 1). In addition, we crossed the *mus81* mutation into *pif1 brca2, pol32 brca2,* and *pif1 pol32 brca2* mutant backgrounds. In all three of these experiments, loss of MUS81 failed to rescue the synthetic lethality (Table 1). Together, these results suggest that the role(s) of PIF1 and POL32 in promoting survival in the absence of BRCA2 may be different from their roles in mammals. Alternatively, replication fork restart in Drosophila may be less dependent on MUS81 than in mammals.

## Discussion

Pif1 family members have overlapping but distinct functions in different organisms and are differentially required for viability. We anticipated that the identification of phenotypes in Drosophila *pif1* mutants might provide additional insight into this gene’s diverse roles in maintaining genomic integrity. Based on our results, it appears that some, but not all, of the functions of PIF1 in other organisms are conserved in Drosophila. For example, Drosophila *pif1* mutant larvae treated with high concentrations of hydroxyurea die prior to adulthood, mimicking the HU sensitivity of both budding and fission yeast. On the other hand, the lack of Drosophila *pif1* mutant sensitivity to many other mutagens suggests that the fly PIF1 does not play a role in the oxidative stress response or in repair of base damage, unlike its yeast orthologs, or that PIF1 has a redundant role in these processes.

### A critical role for Drosophila PIF1 during replication stress

In addition to finding that *pif1* mutants fail to survive exposure to high doses of hydroxyurea, we also identified a vital role for PIF1 during early Drosophila embryogenesis, at a time of extremely rapid genome replication and chromosome segregation. We hypothesize that both phenotypes could be due to failure or impairment of the same mechanism. Eggs produced by *pif1* homozygous mutant mothers rarely hatch and this phenotype cannot be rescued by a wild-type paternal copy of PIF1, indicating that the requirement for PIF1 is most acute during the first two hours of embryo development. The nuclear abnormalities seen in *pif1* mutants during this time are consistent with defects in DNA replication during early embryogenesis. Intriguingly, human PIF1 helicase has been shown to support DNA replication and cell growth during oncogenic stress (Gagou *et al.* 2014). Thus, one conserved feature of PIF1 in all organisms may be to promote efficient DNA synthesis during replication stress.

Based on our findings and studies done in other organisms, we suggest that PIF1 might promote efficient replication during embryogenesis and conditions of replicative stress via one or more of four non-exclusive mechanisms. First, it might work alongside or in combination with POL32 to promote Pol δ processivity. Flies lacking POL32 are also sensitive to HU (our unpublished data) and eggs lacking POL32 have severe nuclear division defects during embryogenesis and fail to develop to larval stages (Tritto *et al.* 2015). In addition, *pol32* mutant embryonic defects cannot be rescued by paternal expression of POL32 (Tritto *et al.* 2015), similar to the *pif1* mutant phenotype.

Second, Drosophila PIF1 might be needed for resolution of G4 or other DNA secondary structures during replication. Although, we have no direct evidence showing that Drosophila PIF1 unwinds G4 DNA, previous studies with *S. cerevisiae* have revealed that DNA replication through G4 motifs is promoted by the PIF1 helicase, where PIF1 unwinds G4 DNA and keeps it unfolded to prevent replication fork stalling and DNA breakage (Paeschke *et al.* 2011; Zhou *et al.* 2014). In addition, *S. pombe* lacking Pfh1 have unresolved G4 structures that lead to increased fork pausing and DNA damage near G4 motifs (Sabouri *et al.* 2014). Strikingly, both yeast and human Pif1 are much more efficient at unwinding G4 DNA than human WRN or budding yeast Sgs1 (Paeschke *et al.* 2013). Thus, it is possible that PIF1 is the main helicase performing G4 structure resolution and/or removal of secondary DNA structures during early embryonic stages. In its absence, the formation of DNA secondary structures stalls replication forks, leading to an accumulation of unresolvable replication intermediates that results in the chromosome segregation defects observed in *pif1* mutant embryos.

Third, PIF1 might be important for completion of replication. Recently, ScPif1 was shown to play a role in replication termination (Deegan *et al.* 2019). Under-replicated DNA intermediates might manifest as fine DNA linkages between segregating chromosomes and chromosome clumps, both of which we observed in *pif1* embryos. These damaged nuclei will be subject to nuclear fallout, resulting in gaps in the typically uniform nuclear monolayer.

A fourth possibility is that PIF1 is needed for cellular responses to replication fork stalling. In budding yeast, the restart of stalled replication forks is dependent on several DNA helicases, including BLM, WRN, FANCM, and PIF1, many of which are recruited to forks by RPA or interact with RPA at forks (Bachrati and Hickson 2008; Luke-Glaser *et al.* 2010; Ramanagoudr-Bhojappa *et al.* 2014). Additionally, fork restart can be mediated by the cleavage of a regressed fork structure, followed by a BIR-like process (Hanada *et al.* 2007; Mayle *et al.* 2015). Since BIR is dependent on Pif1 in *S. cerevisiae*, it is possible that *pif1* embryos are unable to rapidly process stalled replication forks, resulting in slowed replication that manifests as improper syncytial nuclear division and embryonic development.

One of the most surprising results from this study is that *pif1 brca2* and *pol32 brca2* mutants, but not the corresponding *rad51* double mutants, exhibit synthetic lethality at late larval stages. While we showed that PIF1 and POL32 are needed prior to this stage of development, we created homozygous double mutants by mating flies heterozygous for both mutations *in cis*. Thus, maternal deposits of the proteins may have allowed for survival to late larval stages. BRCA2 has been shown to promote replication fork stability in mammalian systems by protecting stalled replication forks from extensive nucleolytic degradation (Schlacher *et al.* 2011; Lemacon *et al.* 2017; Mijic *et al.* 2017). Drosophila BRCA2 has RAD51-independent roles at stalled forks during meiosis (Klovstad *et al.* 2008). Our finding that the *pif1* and *pol32* mutations are lethal in a *brca2*, but not *rad51*, mutant background, suggests that fly BRCA2 may also be important for the stabilization of stalled forks in mitotically dividing cells and supports the idea that it shares a fork protection function with its mammalian counterpart.

We hypothesize that Drosophila PIF1 and POL32 have crucial functions in stabilizing and/or restarting stalled replication forks in the absence of BRCA2. In support of this, human POL32 was shown to mediate the initial DNA synthesis for replication fork restart in the absence of Rad51 (Moriel-Carretero and Aguilera 2010; Mayle *et al.* 2015; Lemacon *et al.* 2017). Specifically, MUS81 cleavage rescues resected forks in BRCA2-deficient cells through a BIR-like mechanism mediated by POL32-dependent DNA synthesis. PIF1 might promote this synthesis, similar to its role in gap repair by HR (see below). However, our failure to observe any rescue of *pif1 brca2* lethality when we removed MUS81 and our successful recovery of viable *pif1 mus81* and *pol32 mus81* mutant adults do not support this model. Alternatively, Pif1 may help with the stabilization of stalled replication forks through pairing of parental ssDNA, migrating HJ structures, and/or directly participating in repair of the lesion (George *et al.* 2009; Ramanagoudr-Bhojappa *et al.* 2014).

### *Drosophila* PIF1 promotes long-range synthesis during HR

Using a quantitative assay of double-strand gap repair in the male pre-meiotic germline, we found that lack of POL32, but not PIF1, results in decreased processive repair synthesis and a significant increase in aborted HR repair. However, a requirement for PIF1 was observed in the absence of POL32. In budding yeast, Pif1 is required for efficient BIR, where it promotes D-loop migration during Pol δ-mediated repair synthesis. A recent study demonstrated that PIF1 may also be important during BIR in Drosophila (Bhandari *et al.* 2019). It is possible that the creation of a 14-kb double-strand gap after *P*-element excision induces a BIR-like mechanism of repair, where the two ends of the break invade homologous templates independently of each other and synthesis proceeds via migrating bubble structures. Supporting this idea, repair of a gap greater than 3 kb depends on POL32-mediated BIR in yeast (Jain *et al.* 2009).

Previous studies of double-strand gap repair in Drosophila led to the model that repair of large gaps by two-ended synthesis-dependent strand annealing requires multiple cycles of strand invasion, synthesis, and displacement of the nascent strand (Mcvey *et al.* 2004). Genetic evidence suggested that translesion polymerases Pol η and Pol ζ might be utilized during initial synthesis, with a subsequent switch to Pol δ that results in more processive synthesis (Kane *et al.* 2012). This model was supported by the observation that loss of both POL32 and the catalytic subunit of Pol ζ resulted in less repair synthesis than was observed in either single mutant. Here, we have shown that *pif1 rev3* mutants behave differently from *pol32 rev3* mutants, consistent with the idea that PIF1 only becomes important during gap repair when Pol δ is impaired by mutation of POL32.

One speculative model, consistent with all these data, is that POL32 acts in the context of the Pol δ complex to promote processivity, while PIF1 is important for the progression of the mobile D-loop. PIF1 could act in concert with PCNA to promote Pol δ-mediated strand displacement at the front of the replication bubble (Buzovetsky *et al.* 2017), and/or behind the D-loop to unwind the newly-synthesized strand to relieve topological hindrance of the nascent strand with the template strand (Wilson *et al.* 2013). If POL32 is present, then Pol δ can engage in strand displacement synthesis and the requirement for PIF1 activity is removed.

### Summary

Collectively, our data are consistent with multiple roles for Drosophila PIF1. First, it is needed to deal with both endogenous and exogenous replication stress. During early embryogenesis, PIF1 acts to ensure proper DNA replication and subsequent chromosome segregation. In addition, it promotes survival when replication forks are challenged by hydroxyurea. Second, in the absence of BRCA2, PIF1 is needed for development past late larval stages, a function that it shares with POL32. Because HR-defective *pif1 rad51* and *pol32 rad51* mutants survive to adulthood, PIF1 and POL32 are likely acting to protect regressed forks or promote replication restart. Third, PIF1 promotes extensive DNA repair synthesis during homologous recombination repair of double-strand gaps in a POL32-independent manner. Together, these findings suggest that Drosophila PIF1 shares some, but not all, of the roles filled by its yeast and mammalian counterparts and lays the groundwork for future investigations into additional PIF1 functions in metazoans.

## Acknowledgments

We thank Jeff Sekelsky, Jim Haber, Catherine Freudenreich, and Kent Golic for helpful discussions, Varandt Khodaverdian for technical help, and the Bloomington Drosophila Stock Center for fly stocks. This research was funded by grants P01GM105473 from the NIH and MCB1716039 from the NSF.

