## Supplemental Figures for "The *Drosophila melanogaster* PIF1 helicase promotes survival during replication stress and processive DNA synthesis during double-strand gap repair"

**Kocak *et al*.**

**Supplemental Figures 1-5**

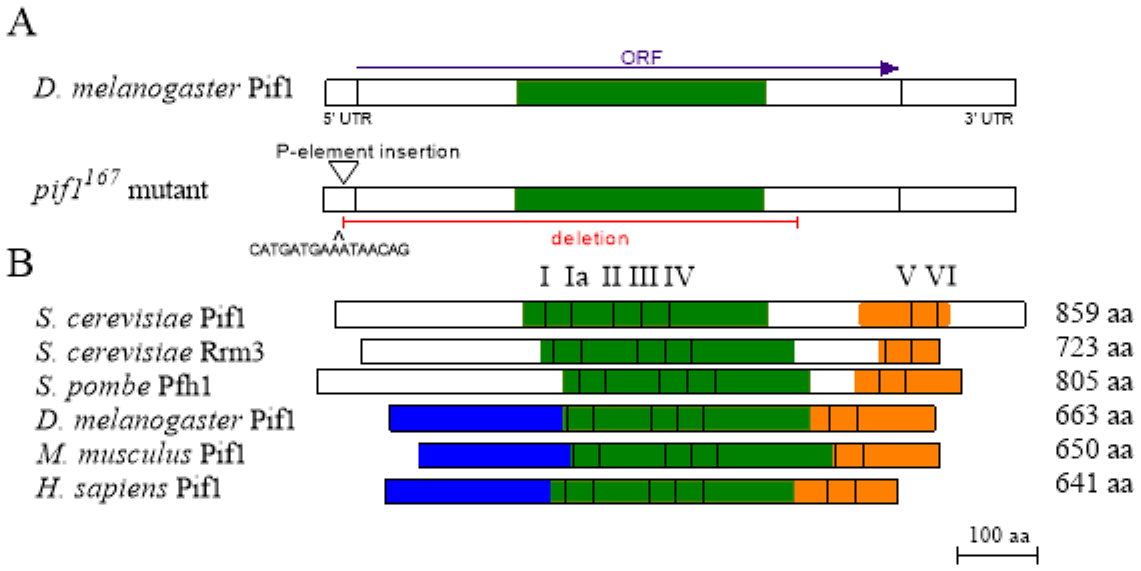


**Figure S1 –** Creation of a Drosophila *pif1* mutant. (A) A large deletion that removes most of the *PIF1* coding sequence was created through imprecise excision of a *P* element inserted in the 5' UTR. The extent of the deletion is indicated by a red bar. A carat indicates an accompanying insertion at the deletion site. (B) Alignment of PIF1 orthologs from multiple species. The conserved helicase domain is shown in green, the conserved N-terminus in shown in blue, and the conserved C-terminal is shown in orange. The seven conserved helicase motifs are indicated in roman numerals and represented by black bars.

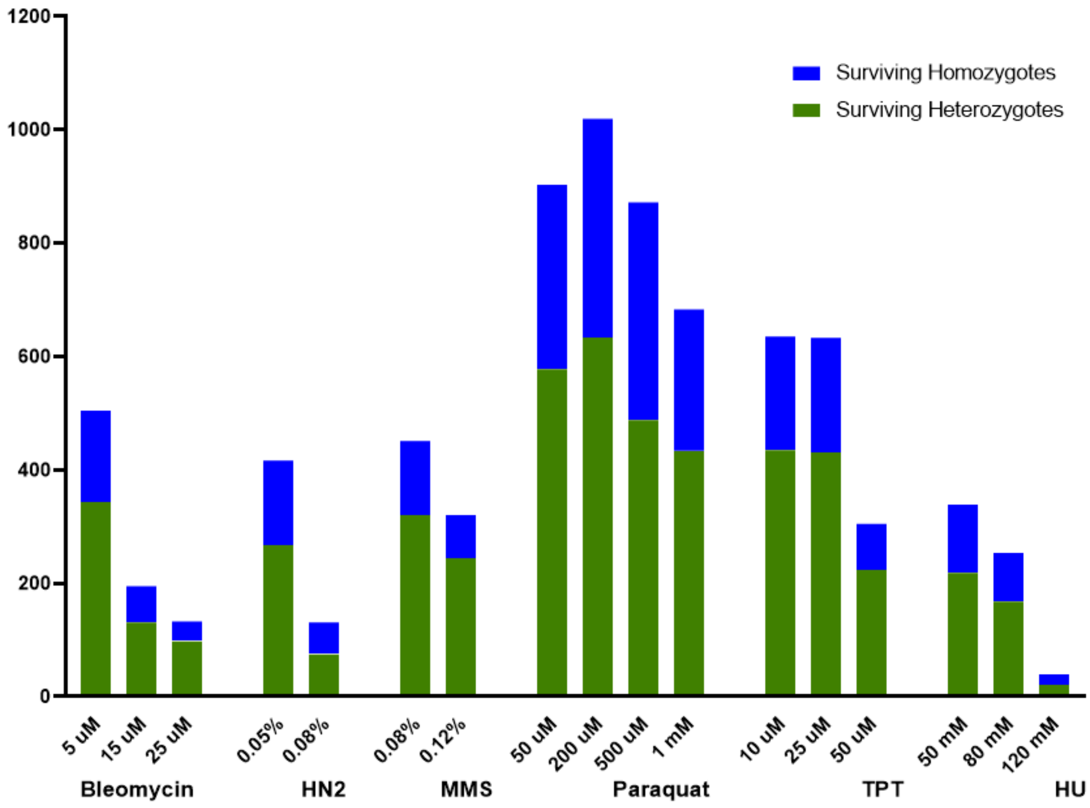


**Figure S2** – Numbers of surviving adults (both *pif1* heterozygotes and homozygotes) for increasing doses of bleomycin, nitrogen mustard (HN2), methyl methane sulfonate (MMS), paraquat, topotecan (TPT) and hydroxyurea (HU).

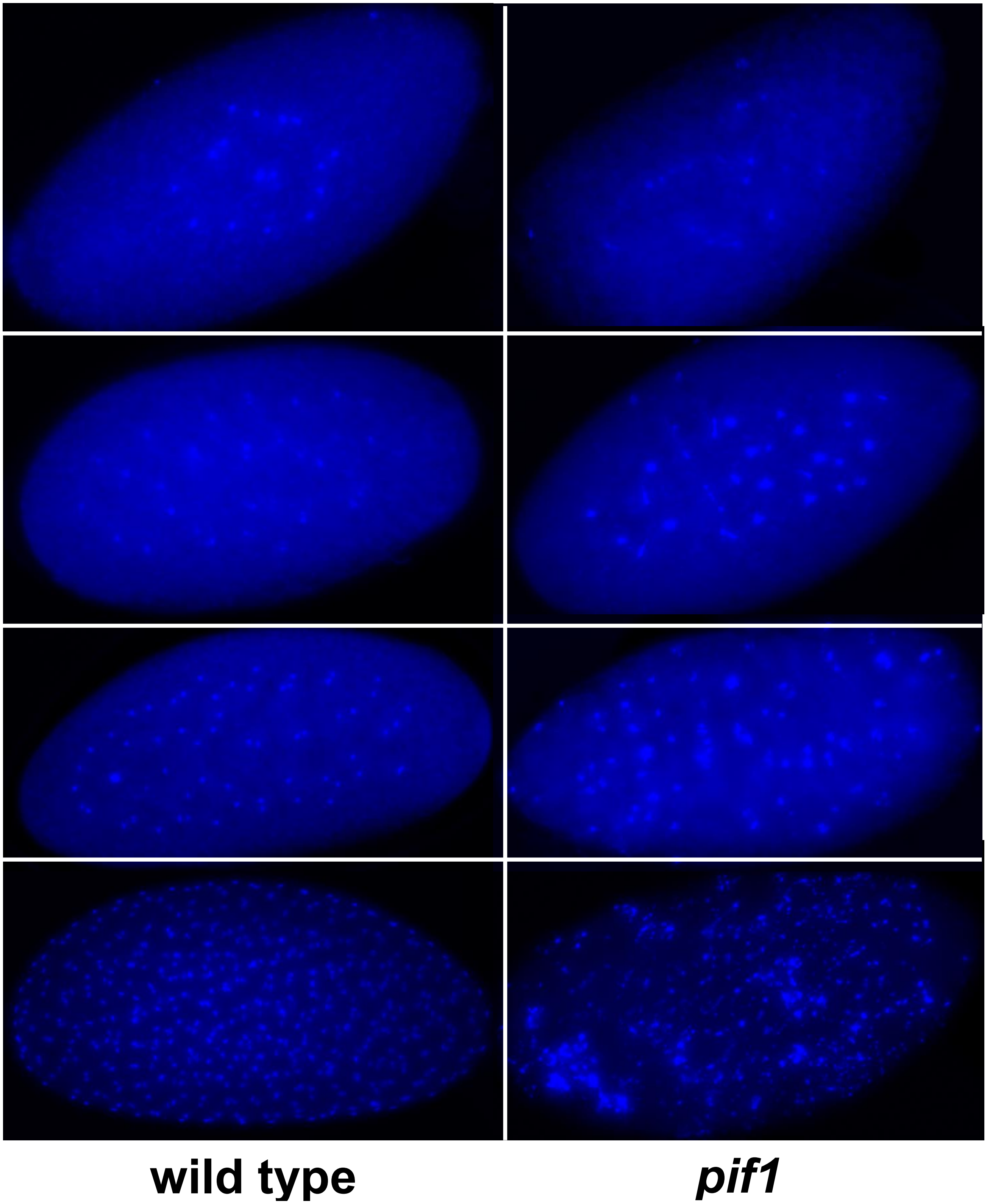


**Figure S3**: DAPI-stained embryos, showing defects in *pif1* mutants occur very early in embryogenesis.

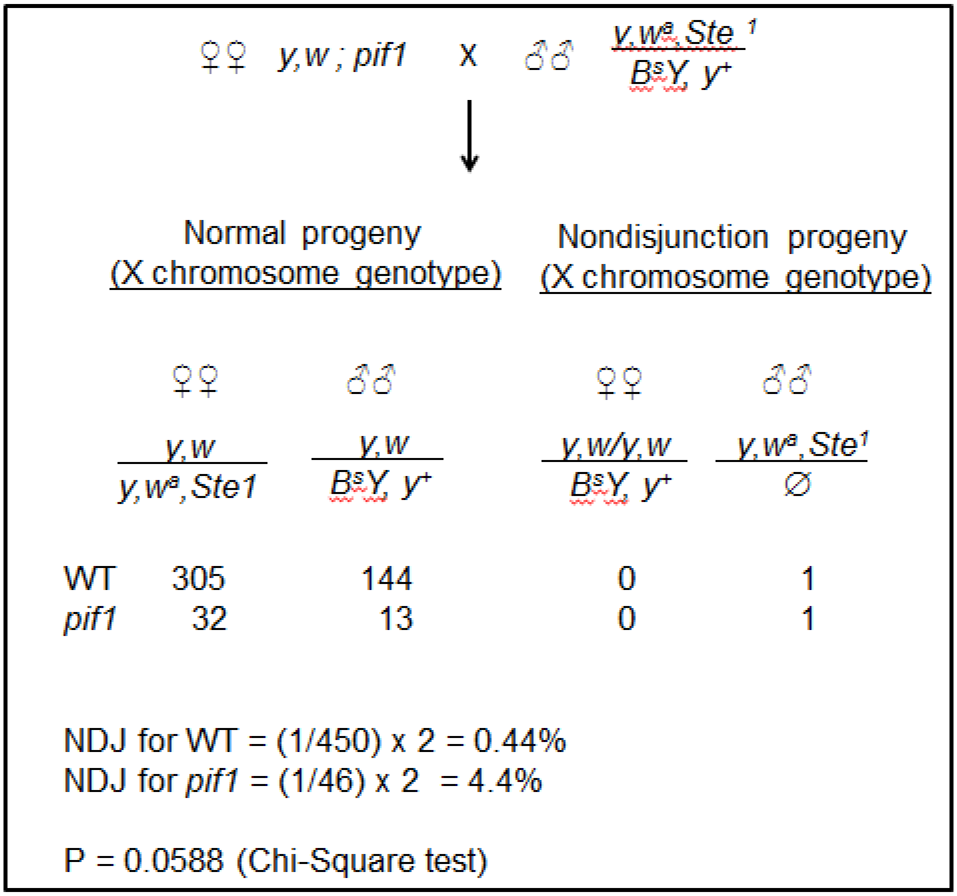


**Figure S4**: *pif1* mutants have a non-significant increase in *X* chromosome nondisjunction. Wild type (WT) or *pif1* mutant females were mated to males of the indicated genotype. Gametes from females in which *X* chromosome nondisjunction occurred will be *yw/yw* or nullo-X. Fertilization of the *yw/yw* egg by a Y-bearing sperm will result in females with Bar-shaped eyes and wild-type body color. Fertilization of the nullo-X egg by an X-bearing sperm will result in males with normal-shaped eyes and yellow bodies. The frequency of non-disjunction is calculated by dividing the number of nondisjunction progeny by the total number of progeny and multiplying by 2 to account for the lethal combinations.

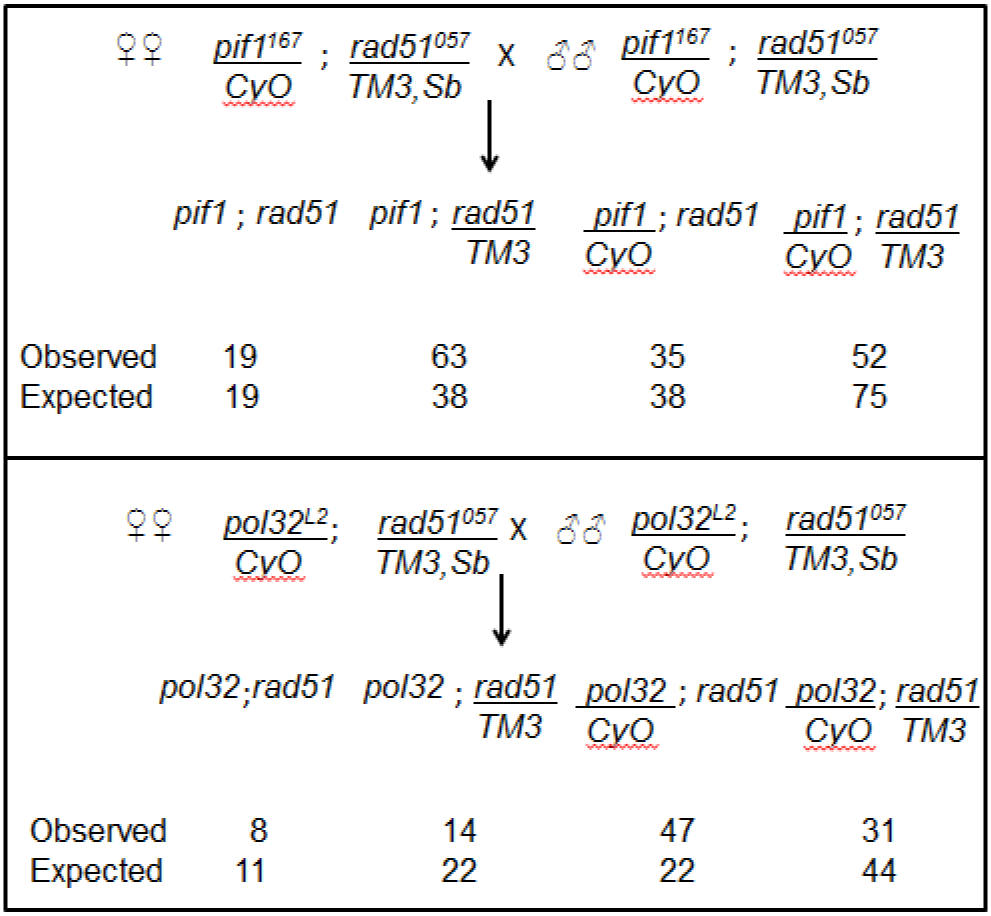


**Figure S5**: Mutation of RAD51 is not lethal in a *pif1* or *pol32* mutant background. Females and males heterozygous for the indicated mutations were mated and the progeny that survived to adulthood were recorded.
